# Physics-Diffusion-Driven Multiscale Aggregation for Drug-Target Interaction Prediction

**DOI:** 10.1101/2025.11.03.686185

**Authors:** Jiongxin Liu, Jiameng Le, Chuanru Wei, Mingming Liu, Zixuan Yin, Yongnan Luo, Hong Qin, Guangbo Yu

## Abstract

Drug-target interaction prediction is an important task in computational drug discovery. To address the limitations of existing graph neural network approaches in propagation depth and robustness, we propose PDDMA-DTI (Physics-Diffusion-Driven Multiscale Aggregation for Drug-Target Interaction Prediction), a physics-inspired, multiscale adaptive aggregation framework. PDDMA-DTI injects a physics-motivated diffusion operator into a heterogeneous drug-target network: global diffusion models cross-modal information propagation via the graph Laplacian, while local diffusion performs neighborhood smoothing in the low-dimensional embedding space. Both processes are solved with the implicit Euler scheme to ensure numerical stability. For each node, we introduce an adaptive stopping rule based on steady-state convergence and a weighted fusion strategy. This strategy leverages hop counts and node degrees to aggregate multi-step diffusion snapshots into a final representation that preserves original features while capturing multi-scale context. To emphasize strong interaction pathways, we further propose a Physical Interaction Summation Enhancement module that augments pairwise interaction features with a linear plus quadratic energy term, thereby amplifying highly correlated drug-target pairs. Extensive experiments on multiple benchmark datasets show that PDDMA-DTI consistently outperforms representative baselines in accuracy and robustness.

## 1 Introduction

A drug target is a biological macromolecule that can bind to a drug and elicit a specific biological function [1]. Identifying drug-target interactions (DTIs) is important for drug repositioning and novel drug discovery. While experimental methods, such as high-throughput screening and in vitro binding assays, are reliable, they are often time-consuming, low-throughput, and costly [2]. With the advancement of deep learning in life sciences, computational prediction based on large-scale databases has become an effective approach to reduce costs and accelerate drug screening [3]. Deep learning-based DTI prediction methods can be broadly categorized into two classes. The first class is based on sequence or structural features: these methods use the SMILES representation of drugs [4] and the amino acid sequences of targets [5] to extract representations for drugs and targets separately, followed by matching for prediction. For instance, Öztürk et al. [6] used two convolutional neural networks [7] to extract features from drug SMILES and target sequences, respectively, to predict drug-target affinity. The second class of methods constructs homogeneous or heterogeneous biological networks from DTIs, drug-drug interactions (DDIs), target-target interactions (TTIs), etc., and utilizes graph models or graph neural networks [7] to mine latent relationships. For example, Yamanishi et al. [8] integrated chemical space, genomic space, and the DTI bipartite graph into a “pharmacological space” and inferred unknown interactions using graph theory.

Building upon this, several studies have extended model architectures or input modalities. For instance, iGRLDTI aims to construct a DTI-centric heterogeneous network and leverages network learning of node relationships to aid prediction. HMSA-DTI [10] introduces a hierarchical, multi-channel self-attention mechanism to achieve crossmodal deep fusion. Meanwhile, NASNet-DTI [11] incorporates Neural Architecture Search into DTI to automatically discover network structures suited to the task. Although these methods have made progress in various aspects, several problems remain inadequately addressed: most methods still rely on globally fixed propagation depths or uniform fusion strategies, making it difficult to adaptively balance “local noise” and “global information” at the node level; some schemes based on deep stacking or strong smoothing regularization, while increasing the propagation range, easily introduce over-smoothing or noise accumulation. To address the aforementioned issues, we propose a PhysicsDiffusion-Driven Multiscale Aggregation framework for Drug-Target Interaction Prediction. Compared with existing approaches, the main contributions of PDDMA-DTI are:

1. Multi-level physics-informed diffusion modeling: We formulate a two-level diffusion scheme—global and local. Global diffusion leverages the graph Laplacian to model cross-modal information propagation and preserve topological consistency, while local diffusion performs neighborhood smoothing in the low-dimensional embedding space to enforce local prediction coherence. Both processes are solved with the implicit Euler method to ensure numerical stability and scalability.
2. Structure-Aware Multi-Scale Fusion Strategy: We introduce a novel weighting rule that explicitly incorporates hop count and node degree to adaptively fuse intermediate representations from multiple diffusion steps. This strategy dynamically balances the contributions of different propagation depths: smaller hop counts receive higher weights to preserve local information, while nodes with higher degrees are assigned greater importance due to their structural reliability.
3. Physical interaction summation enhancement: We propose a physics-motivated interaction enhancement module that models drug-target pairwise interaction energy using a linear plus quadratic term. This mechanism amplifies highly correlated pairs and accentuates strong interaction pathways, thereby improving the model’s ability to distinguish true DTIs.

## 2 Overview of PDDMA-DTI

The proposed PDDMA-DTI model integrates physics-inspired graph enhancement and diffusion learning mechanisms to achieve robust and accurate drug-target interaction prediction. As illustrated in Figure 1, the overall framework consists of four core modules: (A) Drug-Target Network Construction, (B) Physically Guided Diffusion Aggregation, (C) Physical Interaction Summation Enhancement, and (D) DTI Prediction.

**Fig. 1:**
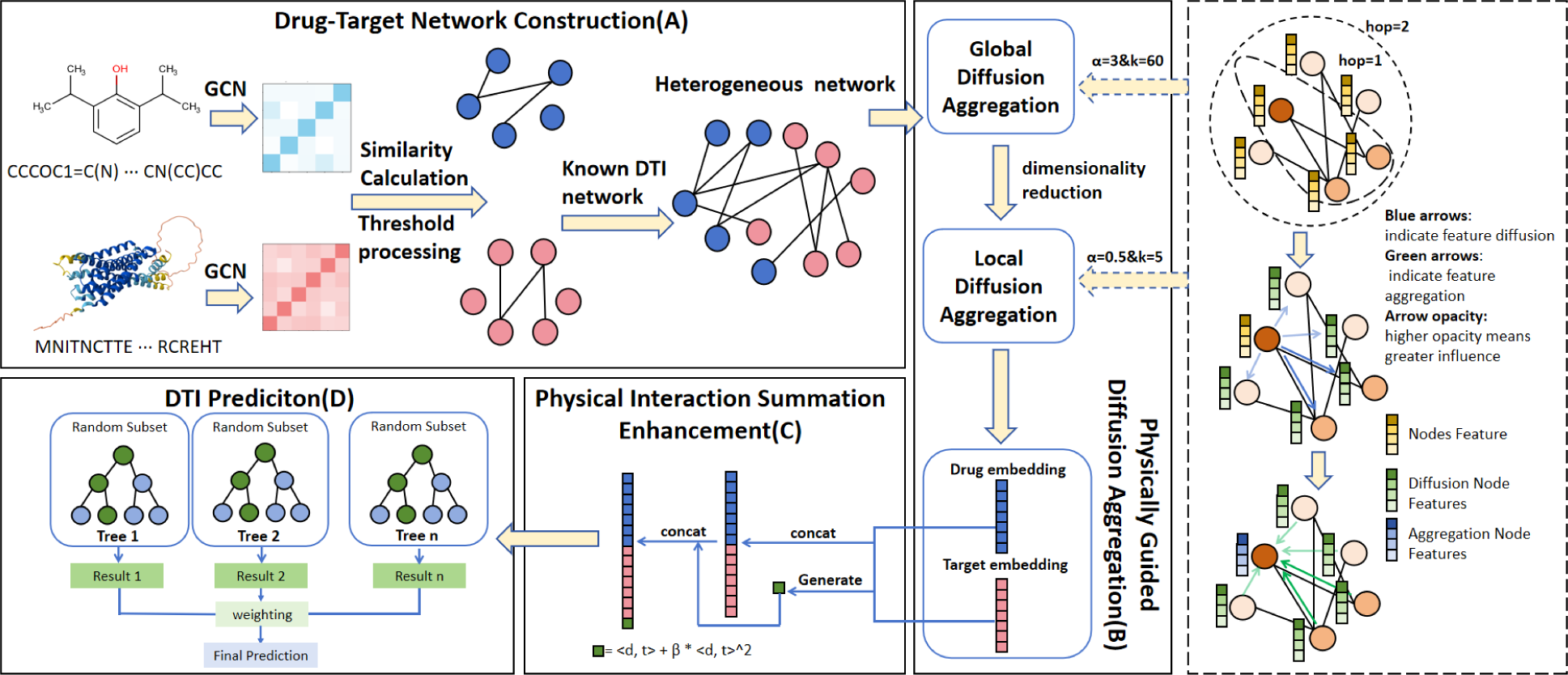
Overview of PDDMA-DTI framework. **Drug-Target Network Construction(A)**: atomic-level drug encoding and protein sequence graph construction; **Physically Guided Diffusion Aggregation(B)**: global and local diffusion with structure-aware fusion; **Physical Interaction Summation Enhancement(C)**: energy-based interaction amplification; **DTI Prediction(D)**: ensemble classification with GBDT.

The Drug-Target Network Construction module begins by encoding each drug from its SMILES string into a molecular graph, from which structural features are extracted using a Graph Convolutional Network(GCN)[12]. In parallel, each protein sequence is represented as a graph of consecutive residues and passed through a two-layer GCN to generate its embedding. To define the graph topology, biologically relevant intra-domain edges are first identified by thresholding the drug-drug and protein-protein similarity matrices. These inferred connections are then combined with known drug-target interactions to form a heterogeneous adjacency matrix that integrates both structural and functional associations. The Physically Guided Diffusion Aggregation module employs hierarchical diffusion combining global and local propagation. The global diffusion algorithm achieves topological consistency via the graph Laplacian, while the local diffusion algorithm performs neighborhood smoothing. Both global and local diffusion utilize a structure-aware fusion strategy based on hop count and node degree. The Physical Interaction Summation Enhancement module encodes pairwise interaction energy through linear-quadratic formalism, using quadratic terms to selectively enhance strongly coupled drug-target pairs and sharpen key interaction pathways. Finally, the DTI Prediction module employs Gradient Boosting Decision Trees to integrate enriched features, capturing complex nonlinear relationships for accurate interaction prediction.

## 3 Methodology

### 3.1 Node Feature Extraction

To capture the detailed chemical structure and topological properties of drug molecules, we utilized the RDKit library [13] to perform per-atom encoding of the SMILES representations. This encoding integrates various atomic attributes, including atom type (e.g., C, N, O, etc., covering 43 common types plus an unknown type), atomic degree, number of explicit/implicit hydrogens, implicit valence, and whether the atom is aromatic. These attributes are converted into high-dimensional binary vectors via one-hot or one-of-k encoding. The atom features are then aggregated through a Graph Convolutional Network to obtain the drug embedding:

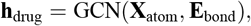

where 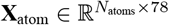 denotes the atom feature matrix, and **E**_bond_ represents the edge indices for bond connections. For protein targets, we first employ one-hot encoding on the amino acid sequence, mapping each amino acid to a 22-dimensional vector (covering the 20 standard amino acids plus ‘X’ and ‘U’) to capture the biochemical diversity of the sequence. A sequence graph is then constructed, where adjacent residues are connected by edges, forming a linear chain-like structure. The protein embedding is extracted via a two-layer Graph Convolutional Network:

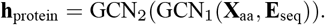

It captures local patterns in the sequence and global context, effectively transforming sequence information into a low-dimensional vector representation suitable for subsequent similarity assessment, graph fusion, and interaction prediction.

### 3.2 Feature Similarity Calculation

To precisely quantify the structural similarity and functional relevance between drugs, we combine two complementary fingerprint representations: Morgan fingerprints [14](a circular fingerprint used to capture local substructures and topological patterns in molecules) and MACCS fingerprints [15](a fingerprint based on predefined structural keys, used to identify key functional groups and pharmacologically relevant features in molecules). The similarity is calculated using a weighted Tanimoto coefficient:

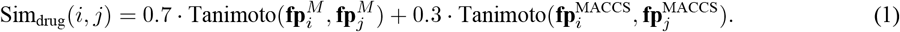

Here, Sim_drug_(*i, j*) denotes the comprehensive similarity between drug *i* and drug *j*, Tanimoto (·, ·) is the Tanimoto similarity coefficient function, 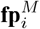, 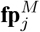 are the Morgan fingerprint vectors for drugs *i* and *j*, and 0.7 and 0.3 are the weight coefficients for the two fingerprint types. The weight assignment emphasizes the structural sensitivity of the Morgan fingerprint and the functional orientation of the MACCS fingerprint. Edges are established by filtering with a high threshold, ensuring that only highly similar drug pairs are connected, thereby constructing a sparse yet informative drug-drug edge set. For protein similarity calculation, this study adopts the Cosine Similarity [16] to measure the similarity between protein embedding vectors. Its core idea is to measure similarity by calculating the cosine of the angle between two vectors, defined as:

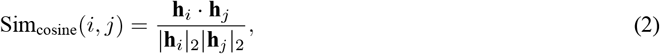

where **h**_*i*_ and **h**_*j*_ represent the feature embedding vectors of protein *i* and *j*, respectively, · denotes the vector dot product, and | · | _2_ denotes the ℓ_2_ norm. Its value range is [−1, 1]. A value close to 1 indicates that the two vectors are highly aligned in direction, meaning the corresponding proteins have high functional or structural similarity in the embedding space; a value close to 0 indicates unrelatedness or orthogonality; a value close to −1 indicates that they are inversely related in semantics or function.

### 3.3 Graph Structure Fusion

The DTI edges are directly obtained from the dataset, representing experimentally verified drug-target associations and forming the core connections of the heterogeneous network. The drug-drug similarity matrix is derived from drugdrug edges, and the target-target similarity matrix is derived from protein-protein edges. The drug-drug similarity matrix, the drug-target matrix, and the target-target similarity matrix are merged into a heterogeneous adjacency matrix containing only 0s and 1s, where 0 indicates no relationship and 1 indicates the existence of a relationship. This fusion strategy integrates multi-source information, promoting global propagation and cross-domain interactive learning.

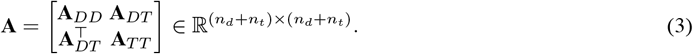

### 3.4 Physically Guided Diffusion Aggregation

To establish a biologically grounded and mathematically rigorous framework for information propagation in drugtarget networks, we introduce a comprehensive physics-informed diffusion paradigm. This approach transforms conventional graph propagation into a physically meaningful process inspired by thermodynamic principles, where information flows through the network following the fundamental laws of diffusion dynamics. The diffusion process initiates with the construction of the graph Laplacian[17] operator **L** = **I** − **D**^−1/2^**AD**^−1/2^, which serves as the mathematical foundation capturing the network’s topological connectivity. This operator essentially quantifies how information naturally dissipates across the heterogeneous graph structure, analogous to heat conduction in physical systems or molecular diffusion in biological contexts. The continuous evolution of node features follows the partial differential equation:

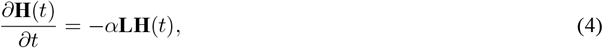

where **H**(*t*) represents the time-dependent node feature matrix, *α* denotes the diffusion coefficient controlling propagation velocity, and the negative Laplacian operator − **L** ensures information flows from high-concentration regions to low-concentration regions, mimicking natural physical processes. For computational implementation on discrete graph structures, we employ the implicit Euler discretization scheme:

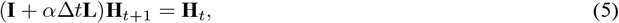

This formulation guarantees unconditional numerical stability regardless of step size, making it particularly suitable for large-scale biological networks where convergence reliability is paramount. The resulting sparse linear system efficiently generates a temporal sequence of diffusion representations {**H**^(0)^, **H**^(1)^, …, **H**^(*K*)^ }, where each **H**^(*k*)^ encapsulates the network’s state after *k* diffusion steps. The three core parameters—diffusion coefficient *α*, maximum step size *K*, and convergence threshold *ϵ*-collectively govern the diffusion behavior. The diffusion coefficient *α* controls propagation velocity: larger values accelerate information flow but risk oversmoothing, while smaller values preserve local features but may limit global information exchange. The maximum step size *K* determines the depth of information propagation: larger *K* enables capture of long-range dependencies but increases computational cost, while smaller *K* focuses on local neighborhoods for efficiency. The convergence threshold *ϵ* balances accuracy and efficiency: smaller *ϵ* ensures thorough diffusion but requires more iterations, while larger *ϵ* accelerates convergence but may terminate propagation prematurely. The global diffusion phase employs aggressive parameters (*α* = 3.0, *K* = 60, *ϵ* = 0.03) to facilitate comprehensive information exchange across the entire heterogeneous network. This configuration allows distant nodes-potentially representing structurally dissimilar drugs or functionally unrelated targets—to influence each other’s representations, thereby capturing long-range dependencies that conventional message-passing might miss.A critical innovation lies in our adaptive termination mechanism, which dynamically determines optimal diffusion depth per node:

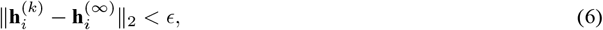

This node-specific convergence checking prevents over-smoothing while ensuring computationally efficient propagation. Nodes in densely connected regions typically converge rapidly, while peripheral nodes require extended diffusion to assimilate global information.The aggregation phase employs a sophisticated multi-scale fusion strategy that intelligently combines diffusion states:

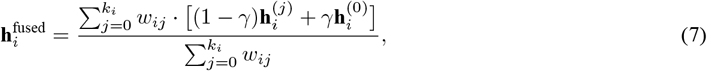

where the adaptive weights *w*_*ij*_ incorporate both temporal decay (favoring recent diffusion steps) and structural importance (prioritizing high-degree nodes), and *γ* balances current diffusion states with original features. This fusion mechanism preserves original node characteristics while integrating progressively broader structural contexts through the diffusion sequence.The resulting fused features **H**^fused^ undergo dimensional projection via a Deep Neural Network:

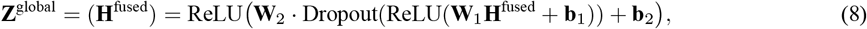

producing compact global embeddings **Z**^global^ ∈ℝ^*N ×*64^ that encode network-wide topological information. Complementing the global perspective, we introduce Local Physics Diffusion to refine embeddings through neighborhoodconstrained propagation:

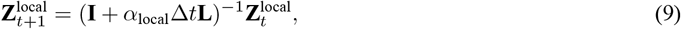

This local diffusion operates directly on the embedding space with conservative parameters (*α*_local_ = 0.5, *K* = 5, *ϵ* = 0.01), emphasizing immediate neighborhood influences while minimizing interference from distant nodes. The small diffusion coefficient (*α*_local_) constrains the propagation intensity, and the limited step count (*K*) confines the influence radius, together ensuring a focused local refinement. This configuration enhances cluster coherence and sharpens decision boundaries between functionally distinct regions. Global diffusion captures system-wide information flow with aggressive parameters, simulating how drug effects or protein functions propagate through the entire interactome. In contrast, local diffusion focuses on community-specific patterns with conservative parameters reflecting localized biological modules or functional complexes. Both processes share the same physical foundation—Laplacian-based diffusion—but operate at different spatial scales with complementary parameterizations. This dual-resolution approach provides unprecedented modeling flexibility, enabling the detection of both broad functional associations and precise interaction specificities.

As illustrated in Figure 2, our adaptive fusion strategy demonstrates significant advantages over conventional approaches. The uniform fusion strategy assigns equal weights to all diffusion steps, failing to capture the varying importance of different propagation depths. The hop-based strategy considers only temporal decay, neglecting structural importance. In contrast, our approach dynamically balances both temporal and structural factors, resulting in more biologically meaningful feature representations. Figure 3 provides a comprehensive 3D visualization of our fusion weight distribution. The weight surface clearly demonstrates the dual dependency on both hop count and node degree, with higher weights concentrated in regions of low hop count and high node degree.

**Fig. 2:**
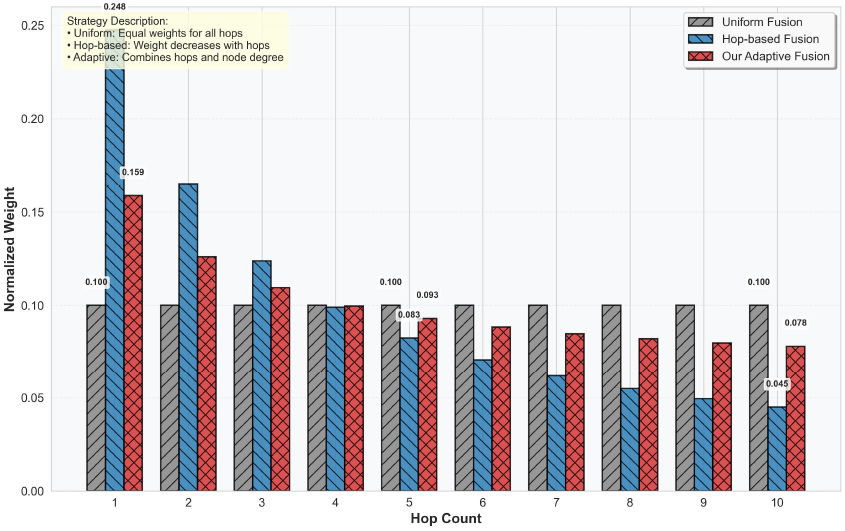
Fusion strategy comparison

**Fig. 3:**
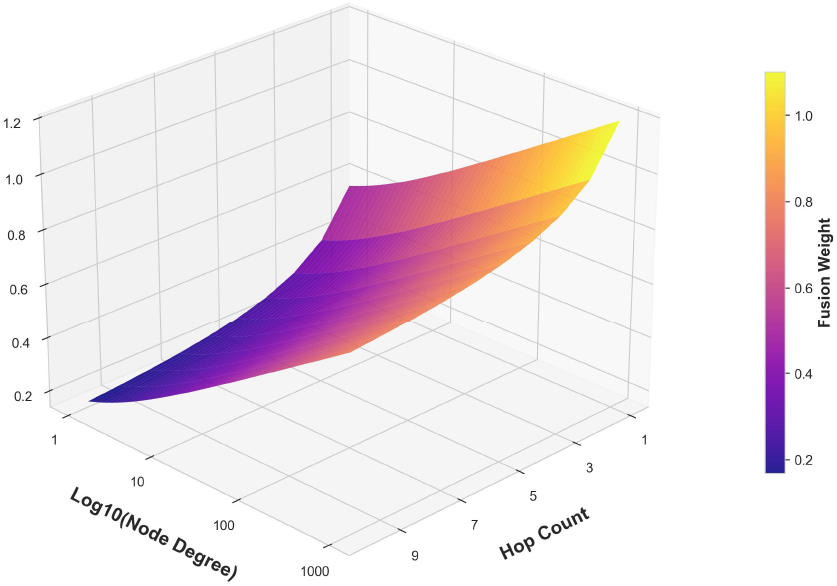
3D weight visualization

### 3.5 Physical Interaction Summation Enhancement

Building upon the physics-informed embeddings, we introduce a novel interaction modeling technique that explicitly quantifies drug-target binding affinities through energy-based formulations. This module translates the abstract embedding similarities into physically meaningful interaction strengths, bridging the gap between topological representation and functional prediction.For each drug-target pair (*d*_*i*_, *t*_*j*_), we define their interaction potential through a composite energy function:

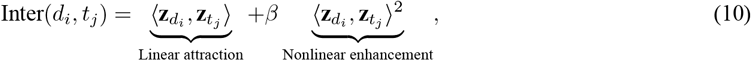

The Physical Interaction Summation Enhancement explicitly models drug-target binding affinities through energy-based computations. The linear term ⟨**z***d*_*i*_, **z***t*_*j*_⟩ captures fundamental complementary patterns between drug and target embeddings, representing basic binding affinity similar to molecular lock-and-key matching. The quadratic term *β* ⟨**z***d*_*i*_, **z***t*_*j*_⟩ ^2^ introduces controlled nonlinear amplification (*β* = 0.01), specifically enhancing strong interaction signals to model cooperative binding effects and allosteric enhancements observed in pharmacological systems. This formulation ensures that highly compatible drug-target pairs receive amplified signals, significantly improving the classifier’s ability to distinguish potent interactions from weak associations. The computed interaction values serve as explicit affinity features that complement the topological information in the concatenated embeddings, providing additional discriminative power for accurate DTI prediction.

### 3.6 Gradient Boosting Decision Tree Classifier

We use XGBoost [18] to classify drug-target pairs. It is an efficient gradient boosting decision tree framework that builds a strong classifier by integrating multiple weak decision tree learners, handling complex nonlinear relationships, and improving prediction accuracy and generalization ability. We first perform a vector concatenation operation, combining the embedding vectors of the drug and the target to form a joint feature representation for the drug-target pair, as shown below.

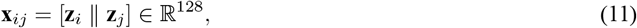

where **x**_*ij*_ is the concatenated feature vector for the drug-target pair (*i, j*), **z**_*i*_, **z**_*j*_ are the node embedding vectors of drug *i* and target *j*, and [· ∥ ·] denotes concatenation. This joint feature representation of the drug-target pair is then fed into XGBoost. The core formula is as follows.

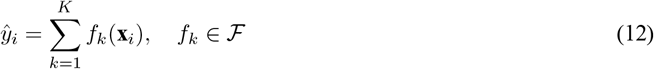

Here, ŷ_*i*_ is the predicted value for sample *i, K* is the total number of trees, *f*_*k*_ is the *k*-th decision tree, **x**_*i*_ is the feature vector of sample *i*, and ℱ is the space of all possible CART tree functions. XGBoost uses an additive model, summing the predictions of multiple weak learners (decision trees) to obtain the final prediction.

## 4 Datasets and Experimental Setup

Experiments were conducted under the “warm-start” scenario using both balanced (1:1 positive-to-negative ratio) and imbalanced (1:10) datasets. The imbalanced setting better reflects real-world conditions where positive samples are typically scarce, while balanced evaluation helps mitigate prediction bias toward negative samples. To minimize data variability, ten-fold cross-validation was employed. The complete dataset was randomly split into 10 subsets, with each subset serving as the test set once while the remaining nine were used for training. Performance was measured using AUROC, AUPR, F1-score, Accuracy, Recall, Specificity, and Precision metrics, implemented with PyTorch. Comprehensive validation was performed on four benchmark datasets (Human[19], BioSNAP[20], BindingDB[21], and DrugBank[22]). For specific experiments, BioSNAP was primarily selected due to its moderate size and comprehensive information. All experiments were run multiple times with optimized parameters, reporting average metrics and standard deviations. Dataset statistics are shown in Table 1.

**Table 1:** Dataset Statistics.

| Dataset | Drugs | Proteins | Positive | Negative |
| --- | --- | --- | --- | --- |
| Human | 852 | 1052 | 3364 | 3387 |
| BindingDB | 14643 | 2623 | 20674 | 28525 |
| DrugBank | 6645 | 4254 | 17511 | 17511 |
| BioSNAP | 4504 | 2181 | 13830 | 13634 |

In our study, we employed four benchmark datasets to evaluate the proposed method. The DrugBank dataset was adopted from a previous study [10], comprising 6,645 drugs and 4,254 targets, with a total of 17,511 drug-target interaction (DTI) edges. The BioSNAP dataset, sourced from prior work [23], includes 4,504 drugs and 2,181 targets, forming 13,830 DTI edges. The BindingDB dataset, originally developed and maintained by the research group of Michael K. Gilson at UCSD, was also obtained from [23]; it consists of 14,643 drugs and 2,623 targets, totaling 20,674 DTI edges. Additionally, the Human dataset, a well-balanced dataset curated by Liu et al. [19], contains 852 drugs and 1,052 targets.

## 5 Experimental Results

### 5.1 Overall Experimental Setup

For comparison, we use the following competitive methods for Drug-Target Interaction prediction as baselines:

- **iGRLDTI**[9]: This method predicts DTIs by constructing a DTI-centric heterogeneous network. However, its main limitations lie in the insufficient utilization of drug-drug and target-target relationships, and the neglect of target structural features during feature extraction, which constrains its ability to model complex biological relationships.
- **HMSA-DTI**[10]: While this hierarchical multimodal self-attention GNN framework can capture complex drugtarget interactions through multimodal feature fusion, it primarily focuses on local structural features and exhibits limitations in global topological relationship modeling, with insufficient exploitation of graph structural information.
- **NASNet-DTI**[11]: This method employs a node adaptive learning strategy to dynamically determine aggregation depth, effectively mitigating over-smoothing. However, its influence vector convergence-based adaptive mechanism shows deficiencies in capturing multi-scale structural information, and the simple averaging fusion strategy may not adequately preserve discriminative features from different propagation depths.

Table 2 presents the performance comparison results on balanced datasets. PDDMA-DTI demonstrates significant advantages across all evaluation metrics and datasets. Particularly, the near-perfect performance achieved on the BindingDB dataset (AUC: 0.999, AUPR: 0.999) validates the framework’s exceptional capability in handling large-scale biological network data.

**Table 2:** Model performance comparison on four benchmark datasets (Positive : Negative = 1 : 1).

| Dataset | Method | AUC | AUPR | F1 | Sensitivity | Specificity |
| --- | --- | --- | --- | --- | --- | --- |
| DrugBank | iGRLDTI | 0.918 ± 0.004 | 0.914 ± 0.005 | 0.860 ± 0.008 | 0.834 ± 0.011 | 0.853 ± 0.009 |
|  | HMSA-DTI | 0.922 ± 0.005 | 0.919 ± 0.005 | 0.833 ± 0.007 | 0.856 ± 0.009 | 0.826 ± 0.007 |
|  | NASNet-DTI | 0.934 ± 0.004 | 0.932 ± 0.004 | 0.846 ± 0.007 | 0.848 ± 0.011 | 0.869 ± 0.009 |
|  | **PDDMA-DTI** | **0.991 ± 0.001** | **0.990 ± 0.001** | **0.951 ± 0.005** | **0.966 ± 0.006** | **0.934 ± 0.006** |
| BioSNAP | iGRLDTI | 0.924 ± 0.004 | 0.927 ± 0.004 | 0.841 ± 0.006 | 0.829 ± 0.007 | 0.868 ± 0.008 |
|  | HMSA-DTI | 0.930 ± 0.004 | 0.937 ± 0.004 | 0.863 ± 0.005 | 0.820 ± 0.007 | 0.847 ± 0.006 |
|  | NASNet-DTI | 0.942 ± 0.004 | 0.941 ± 0.004 | 0.864 ± 0.006 | 0.845 ± 0.008 | 0.882 ± 0.007 |
|  | **PDDMA-DTI** | **0.988 ± 0.001** | **0.988 ± 0.001** | **0.943 ± 0.004** | **0.957 ± 0.005** | **0.927 ± 0.009** |
| BindingDB | iGRLDTI | 0.932 ± 0.004 | 0.933 ± 0.004 | 0.853 ± 0.005 | 0.855 ± 0.005 | 0.859 ± 0.006 |
|  | HMSA-DTI | 0.974 ± 0.004 | 0.924 ± 0.004 | 0.962 ± 0.004 | 0.900 ± 0.006 | 0.908 ± 0.006 |
|  | NASNet-DTI | 0.982 ± 0.003 | 0.981 ± 0.004 | 0.926 ± 0.005 | 0.913 ± 0.006 | 0.934 ± 0.007 |
|  | **PDDMA-DTI** | **0.999 ± 0.000** | **0.999 ± 0.000** | **0.986 ± 0.002** | **0.990 ± 0.002** | **0.981 ± 0.004** |
| Human | iGRLDTI | 0.965 ± 0.004 | 0.962 ± 0.005 | 0.901 ± 0.009 | 0.931 ± 0.014 | 0.866 ± 0.013 |
|  | HMSA-DTI | 0.993 ± 0.000 | 0.994 ± 0.000 | 0.957 ± 0.000 | 0.935 ± 0.000 | **0.980 ± 0.000** |
|  | NASNet-DTI | 0.964 ± 0.000 | 0.963 ± 0.000 | 0.898 ± 0.000 | 0.916 ± 0.000 | 0.876 ± 0.000 |
|  | **PDDMA-DTI** | **0.996 ± 0.002** | **0.996 ± 0.002** | **0.976 ± 0.005** | **0.982 ± 0.007** | 0.969 ± 0.010 |

The imbalanced dataset results presented in Table 3 highlight PDDMA-DTI’s exceptional capability to handle real-world data distribution challenges. In practical drug discovery scenarios, imbalanced data is the norm rather than the exception, making this evaluation particularly meaningful. Our framework demonstrates remarkable resilience to class imbalance, maintaining performance levels that closely approximate those achieved on balanced datasets. The framework’s performance stability across varying imbalance conditions further validates its design principles. Unlike conventional methods that often exhibit significant performance degradation when faced with class imbalance, PDDMA-DTI maintains robust predictive capability, suggesting that the integration of physical principles provides a stabilizing influence that transcends data distribution artifacts. This characteristic is particularly valuable for realworld drug screening applications, where the ratio of potential drug-target interactions to non-interactions is inherently skewed. Moreover, the balanced improvements across both AUC and AUPR metrics indicate that our approach successfully addresses the dual challenges of overall ranking accuracy and minority class identification. The framework’s ability to maintain high F1-scores—reflecting a harmonious balance between precision and recall—further confirms its suitability for practical deployment in computational drug discovery pipelines, where both false positives and false negatives carry significant costs.

**Table 3:** Performance comparison on four benchmark datasets(Positive : Negative = 1 : 10).

| Dataset | Method | AUC | AUPR | F1 |
| --- | --- | --- | --- | --- |
| DrugBank (Imbalanced) | iGRLDTI | 0.899 | 0.682 | 0.593 |
|  | HMSA-DTI | 0.824 | 0.534 | 0.709 |
|  | NASNet-DTI | 0.942 | 0.714 | 0.719 |
|  | **PDDMA-DTI** | **0.992** | **0.942** | **0.857** |
| BindingDB (Imbalanced) | iGRLDTI | 0.976 | 0.877 | 0.771 |
|  | HMSA-DTI | 0.921 | 0.729 | 0.846 |
|  | NASNet-DTI | 0.979 | 0.883 | 0.804 |
|  | **PDDMA-DTI** | **0.999** | **0.994** | **0.964** |
| BioSNAP (Imbalanced) | iGRLDTI | 0.936 | 0.702 | 0.603 |
|  | HMSA-DTI | 0.938 | 0.789 | 0.858 |
|  | NASNet-DTI | 0.942 | 0.754 | 0.702 |
|  | **PDDMA-DTI** | **0.992** | **0.944** | **0.865** |
| Human (Imbalanced) | iGRLDTI | 0.975 | 0.850 | 0.731 |
|  | HMSA-DTI | 0.993 | 0.953 | 0.881 |
|  | NASNet-DTI | 0.970 | 0.836 | 0.719 |
|  | **PDDMA-DTI** | **0.998** | **0.979** | **0.925** |

### 5.2 Ablation Study Results Analysis

To evaluate the contribution of each component in PDDMA-DTI, we conducted an ablation study on the BioSNAP dataset (1:10 positive-to-negative ratio) with four configurations: 1) global module only, 2) global + local module, 3) global module + interaction enhancement, and 4) the full model integrating all three. These modules actually all consist of two parts: one is physical diffusion and the other is adaptive aggregation. As illustrated in Figure 4 and Figure 5, the baseline configuration using only the global module already achieved solid performance. The integration of the local module provided a further enhancement in AUPR, while the most notable improvement was observed after incorporating the physical interaction summation component. The full model, combining all components, attained the highest performance, demonstrating a clear synergistic effect among the modules. All model variants exhibited consistently low variance across runs, confirming the robustness of the proposed approach.

**Fig. 4:**
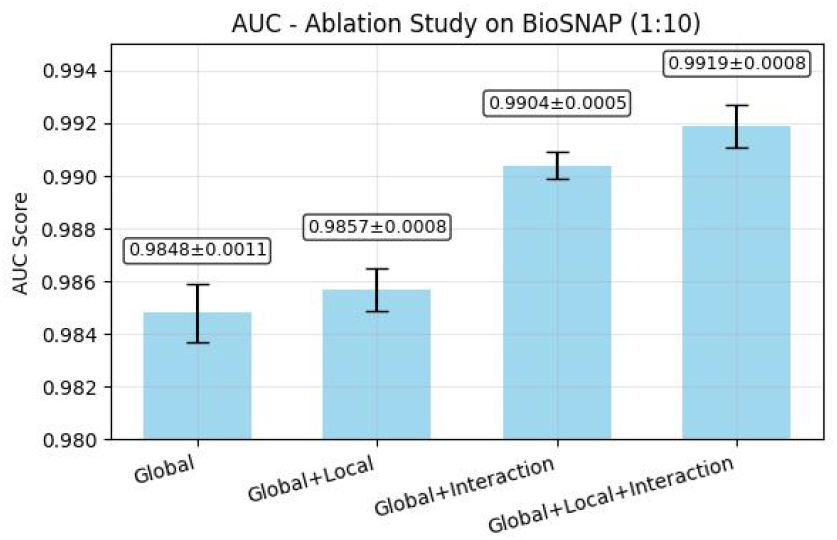
AUC experimental results under different module combinations

**Fig. 5:**
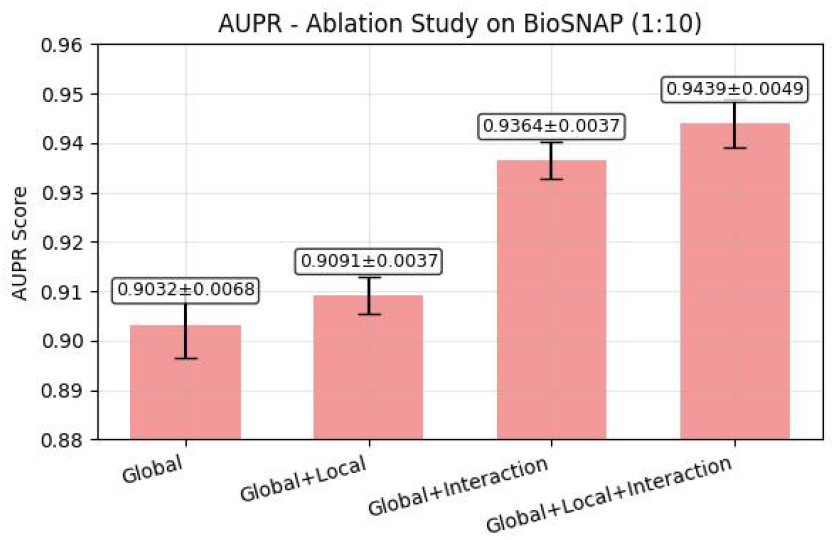
AUPR experimental results under different module combinations

### 5.3 Hyperparameter Sensitivity Analysis

To systematically study the impact of diffusion parameters on model performance, we conducted a grid search and sensitivity analysis on the diffusion coefficient *α* (`diffusion_coef`) and the number of diffusion steps *k* (`physics_steps`).

To exclude interference from other modules, the global/local physics modules and the physical interaction summation enhancement were not enabled in these experiments; only the single physics diffusion and aggregation module was retained as a representative configuration. Relevant results are shown in Table 4 and Table 5, with the main conclusions summarized as follows:

**Table 4:** Impact of different diffusion coefficients on model performance.

| diffusion_coef ($\alpha$) | AUC | AUPR |
| --- | --- | --- |
| 0.1 | 0.8669 | 0.8811 |
| 1 | 0.8875 | 0.8994 |
| 3 | 0.9117 | 0.9183 |
| **5** | **0.9215** | **0.9255** |
| 7 | 0.9203 | 0.9251 |
| 9 | 0.9190 | 0.9242 |
| 15 | 0.9157 | 0.9213 |
| 20 | 0.9181 | 0.9223 |
| 50 | 0.9140 | 0.9199 |

**Table 5:** Impact of different numbers of diffusion steps on model performance.

| physics_steps ($k$) | AUC | AUPR |
| --- | --- | --- |
| 1 | 0.8672 | 0.8810 |
| 2 | 0.9265 | 0.9288 |
| 5 | 0.9564 | 0.9541 |
| 10 | 0.9721 | 0.9699 |
| 20 | 0.9769 | 0.9745 |
| 50 | 0.9770 | 0.9748 |
| **60** | **0.9784** | **0.9764** |
| 70 | 0.5000 | 0.5000 |
| 80 | 0.5000 | 0.5000 |
| 90 | 0.5000 | 0.5000 |

First, with the number of diffusion steps fixed at *k* = 3, we tested the performance changes corresponding to different *α* values. The results show that as the diffusion coefficient increases from 0.1 to 5, the model performance (AUC and AUPR) continuously improves, peaking around *α* = 5 (AUC≈0.92, AUPR≈0.93). When *α* is further increased beyond 15, performance slightly declines, indicating that excessively strong diffusion leads to feature over-smoothing and loss of discriminative information. This result suggests that an appropriate diffusion strength can effectively enhance the feature propagation range and discriminative ability, but excessive diffusion will cause node embeddings to converge, weakening the model’s discriminative power.

Second, with the diffusion coefficient fixed at *α* = 5, we tested the impact of different numbers of diffusion steps *k* on performance. The experiments found that as *k* increases from 1 to 60, model performance improves significantly and reaches the optimum at *k* = 60 (AUC≈0.978, AUPR≈0.976), indicating that moderately increasing the number of diffusion steps helps nodes more fully absorb neighborhood information. However, when the number of steps continues to increase (*k* ≥ 70), performance degrades sharply to a random level (AUC≈0.5), indicating severe feature oversmoothing and numerical degradation. This shows that excessively large values of *k* destroy feature discriminability and numerical stability.

### 5.4 Case Study

We conducted a case study on the BioSNAP dataset to evaluate the method’s capability in identifying novel drugtarget interactions. The experimental procedure was as follows: one drug and one protein were randomly selected as test candidates, and all their original interactions were removed from the dataset. The remaining known DTIs were used to construct the training set, assessing the method’s robustness and generalization ability in a real cold-start scenario.

Specifically, we selected the drug (DrugBank ID: DB00749, Etodolac) and the protein (UniProt ID: P35270, Sepiapterin reductase) as test candidates. For the drug candidate, we constructed 8 positive samples and paired them with 2,172 negative samples. For the protein candidate, we constructed 4 positive samples and 4,500 negative samples. The experimental results are shown in Table 6 and Table 7, presenting the Top-10 predicted rankings for each candidate. After ranking by prediction scores and examining the Top-10 rankings, 2 out of 8 original positive samples for the drug candidate and 3 out of 4 for the protein candidate ranked in the Top-10. These results indicate that our method can effectively identify novel DTIs, demonstrating good generalization capability.

**Table 6:**
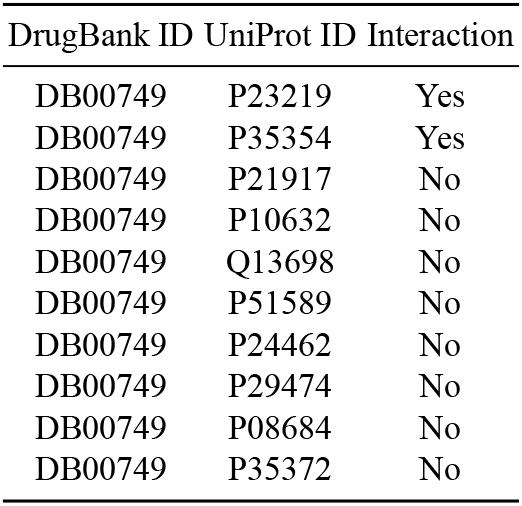
Top-10 Prediction Results for Candidate Drug.

| DrugBank ID | UniProt ID | Interaction |
| --- | --- | --- |
| DB00749 | P23219 | Yes |
| DB00749 | P35354 | Yes |
| DB00749 | P21917 | No |
| DB00749 | P10632 | No |
| DB00749 | Q13698 | No |
| DB00749 | P51589 | No |
| DB00749 | P24462 | No |
| DB00749 | P29474 | No |
| DB00749 | P08684 | No |
| DB00749 | P35372 | No |

**Table 7:**
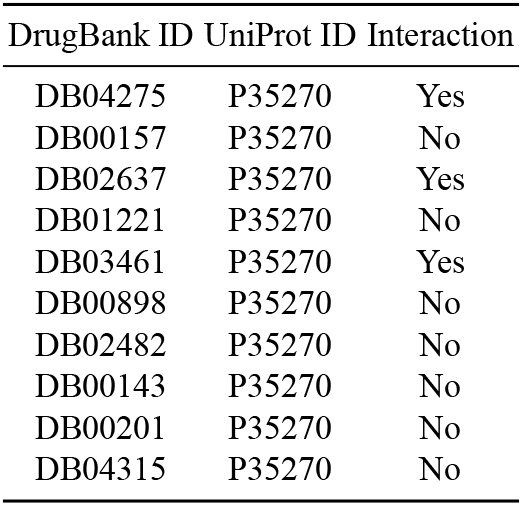
Top-10 Prediction Results for Candidate Protein.

| DrugBank ID | UniProt ID | Interaction |
| --- | --- | --- |
| DB04275 | P35270 | Yes |
| DB00157 | P35270 | No |
| DB02637 | P35270 | Yes |
| DB01221 | P35270 | No |
| DB03461 | P35270 | Yes |
| DB00898 | P35270 | No |
| DB02482 | P35270 | No |
| DB00143 | P35270 | No |
| DB00201 | P35270 | No |
| DB04315 | P35270 | No |

## 6 Conclusion

This paper proposed a DTI prediction framework based on physics-diffusion-driven multiscale aggregation. We constructed a three-level mechanism comprising global physics diffusion and aggregation, local physics diffusion and aggregation, and physical interaction summation enhancement. Numerical stability of the diffusion process is ensured through an implicit Euler solver, and an adaptive fusion strategy based on node-specific convergence steps is designed. Experimental results demonstrate that these innovations collectively enhance both accuracy and robustness.

## 7 Data availability

The code and data are available at “https://github.com/LIUYellowBlack/PDDMA-DTI“

## Acknowledgements

This work was supported by the Tianjin Natural Science Foundation (No.22JCYBJC01020).

## References

[1] Yaping Feng, Qi Wang, and Tao Wang. “Drug target protein-protein interaction networks: a systematic perspective”. In: Biomed Research International 2017 (2017).

[2] J. A. DiMasi, H. G. Grabowski, and R. W. Hansen. “Innovation in the pharmaceutical industry: new estimates of R&D costs”. In: Journal of Health Economics 47 (2016), pp. 20–33.

[3] Jian Peng, Jie Li, and Xuequn Shang. “A learning-based method for drug-target interaction prediction based on feature representation learning and deep neural network”. In: BMC Bioinformatics 21.13 (2020), pp. 1–13.

[4] David Weininger. “SMILES, a chemical language and information system. 1. Introduction to methodology and encoding rules”. In: Journal of Chemical Information and Computer Sciences 28.1 (1988), pp. 31–36.

[5] C. B. Anfinsen. “Principles that govern the folding of protein chains”. In: Science 181.4096 (1973), pp. 223– 230.

[6] H. Öztürk, A. Özgür, and E. Özkirimli. “DeepDTA: deep drug–target binding affinity prediction”. In: Bioinformatics 34.17 (2018), pp. i821–i829.

[7] Franco Scarselli et al. “The Graph Neural Network Model”. In: IEEE Transactions on Neural Networks 20.1 (2009), pp. 61–80.

[8] Yoshihiro Yamanishi, Masaaki Kotera, Minoru Kanehisa, et al. “Drug-target interaction prediction from chemical, genomic and pharmacological data in an integrated framework”. In: Bioinformatics 26.12 (2010), pp. i246– i254.

[9] Bo-Wei Zhao et al. “iGRLDTI: an improved graph representation learning method for predicting drug–target interactions over heterogeneous biological information network”. In: Bioinformatics 39.8 (July 2023), btad451. ISSN: 1367-4811.

[10] Jilong Bian et al. “Hierarchical multimodal self-attention-based graph neural network for DTI prediction”. In: Briefings in Bioinformatics 25.4 (June 2024), bbae293. ISSN: 1477-4054.

[11] Ningyu Zhong and Zhihua Du. “NASNet-DTI: accurate drug–target interaction prediction using heterogeneous graphs and node adaptation”. In: Briefings in Bioinformatics 26.4 (July 2025), bbaf342. ISSN: 1477-4054.

[12] T. N. Kipf and M. Welling. “Semi-supervised classification with graph convolutional networks”. In: Proceedings of the 5th International Conference on Learning Representations. 2017.

[13] A. P. Bento, A. Hersey, E. Félix, et al. “An open source chemical structure curation pipeline using RDKit”. In: Journal of Cheminformatics 12.1 (2020), p. 51. DOI: 10.1186/s13321-020-00456-1.

[14] H. L. Morgan. “The Generation of a Unique Machine Description for Chemical Structures-A Technique Developed at Chemical Abstracts Service”. In: Journal of Chemical Documentation 5.2 (1965), pp. 107–113.

[15] Joseph L. Durant et al. “Reoptimization of MDL Keys for Use in Drug Discovery”. In: Journal of Chemical Information and Computer Sciences 42.6 (2002), pp. 1273–1280.

[16] Ehsaneddin Asgari and Mohammad RK Mofrad. “Continuous distributed representation of biological sequences for deep proteomics and genomics”. In: PloS one 10.11 (2015), e0141287.

[17] R. Merris. “Laplacian matrices of graphs: a survey”. In: Linear Algebra and its Applications 197 (1994), pp. 143– 176. DOI: 10.1016/0024-3795(94)90486-3.

[18] T. Chen and C. Guestrin. “XGBoost: A Scalable Tree Boosting System”. In: Proceedings of the 22nd ACM SIGKDD International Conference on Knowledge Discovery and Data Mining. 2016, pp. 785–794. DOI: 10.1145/2939672.2939785.

[19] Hui Liu et al. “Improving compound–protein interaction prediction by building up highly credible negative samples”. In: Bioinformatics 31.12 (June 2015), pp. i221–i229. ISSN: 1367-4803.

[20] M. Zitnik, R. Sosič, and J. Leskovec. BioSNAP Datasets: Stanford Biomedical Network Dataset Collection. https://snap.stanford.edu/biodata/. 2018.

[21] M. K. Gilson, T. Liu, M. Baitaluk, et al. “BindingDB in 2015: a public database for medicinal chemistry, computational chemistry and systems pharmacology”. In: Nucleic Acids Research 44.D1 (2016), pp. D1045–D1053.

[22] D. S. Wishart, Y. D. Feunang, A. C. Guo, et al. “DrugBank 5.0: a major update to the DrugBank database for 2018”. In: Nucleic Acids Research 46.D1 (2018), pp. D1074–D1082.

[23] P. Bai et al. “Interpretable bilinear attention network with domain adaptation improves drug–target prediction”. In: Nature Machine Intelligence 5 (2023), pp. 126–136.

